# Qing-Luo-Yin eases angiogenesis in adjuvant-induced arthritis rats by activating PPARγ

**DOI:** 10.1101/2024.08.01.606254

**Authors:** Meng-Ke Song, Qin Yin, Meng-Fan Gu, Wen-Gang Chen, Opeyemi Joshua Olatunji, Yan Li, Jian Zuo

## Abstract

**Objective:** Qing-Luo-Yin (QLY) is an anti-rheumatic herbal formula with potentials activating PPARγ. This study investigated if its anti-angiogenesis effects are related to immune modulation.

**Method:** Adjuvant-induced arthritis (AIA) rats were orally treated by QLY or rosiglitazone (a PPARγ agonist) for 30 days. Their immune and metabolism statues were investigated afterward. Isolated monocytes and lymphocytes were co-cultured reciprocally, and treated by different serums. Healthy rats received blood transfusion from QLY-treated or AIA model rats. Two days ahead of sacrifice, a matrigel plug was planted. The plug and some blood immune indicators were examined. AIA rat serum-incubated THP-1 and Jurkat cells were treated by sinomenine, berberine and palmatine. The medium and T0070907 (a PPARγ inhibitor) were used to stimulate HUVEC cells.

**Results:** QLY showed similar therapeutic effects on AIA to rosiglitazone, alleviating joint injuries, synovial angiogenesis, and metabolic disorders. Although QLY impaired inflammatory phenotype of AIA monocytes in vivo, the effect was hardly achieved or sustained in vitro. T cells from QLY-treated AIA rats showed the weakened inflammatory phenotype, and were unable to induce monocytes inflammatory polarization. AIA rat lymphocytes induced angiogenesis in the matrigel plug in healthy recipients. In lymphocytes enrichment site, QLY reduced the secretion of IL-17A, IFNγ, and many angiogenesis-related cytokines. QLY-related components affected Jurkat but not THP-1 cells. Jurkat T cells induced angiogenesis of HUVEC cells when cultured by AIA rat serum. Inhibitory effects of the compounds on it were abolished by T0070907.

**Conclusion:** PPARγ activation in T cells is a foundation for the anti-angiogenesis property of QLY.

## 1. Introduction

The successful treatments of rheumatoid arthritis (RA) are still challenging due to its elusive etiological mechanisms. Many pathological changes contribute to its onset and progression. According to the current understanding, synovitis is the key driving force underlying RA-related structure degradation and functional loss of joints [1-2]. In addition to synovial invasion, angiogenesis fuels this abnormality directly [3]. Because articular manifestations are still the priority of anti-rheumatic therapies now, synoviocytes and vascular endothelial cells should be the main therapeutic targets. Despite many studies have validated plausibility of this strategy [4-5], there is no such targeting drug available now. A fundamental reason is that RA pathogenesis is driven by immune hyper-activation rather than specific local changes [2]. Various outcomes from effective RA therapies can be explained by the improved immune milieu, and immune-regulating approach would bring more benefits than the other strategies.

Thanks to the accumulating knowledge about the immune characteristics of RA, some biological agents have been successfully developed and introduced into clinical practices [6]. Compared to the conventional disease modifying anti-rheumatic drugs (DMARDs), they exhibit advantages in treating various RA-related pathological changes [6]. They mainly target immune cells, the dominant roles in RA pathology, as well as the signals controlling their differentiation. But these drugs are unaffordable in developing countries. Hence, there is still a great demand for cheap and effective therapies.

In addition to DMARDs, herbal medicines are widely used worldwide. They might be even more effective than the synthesized compounds, benefited from their functions on alleviating symptom combinations by numerous chemical combinations and therapeutic targets, which have been explored by novel technologies such as network pharmacology [7-9]. As a successful representative, the Traditional Chinese Medicine (TCM) formula Qing-Luo-Yin (QLY) has been used for over 40 years. Its efficacy has been thoroughly validated by both clinical and pharmacological researches [10-13]. Of note, it exhibited eyes catching effects on synovial angiogenesis [14].

Great endeavor is devoted to clarify the anti-rheumatic mechanisms of QLY, and we confirm that SIRT1/PPARγ is one of its molecular targets [12-13]. Interestingly, SIRT1 and PPARγ function as a pair of rivals, from both the immune and metabolism perspectives. Furthermore, they negatively regulate each other’s expression [15-16]. QLY shows potentials in inhibiting SIRT1 and activating PPARγ [13]. Since SIRT1 is generally recognized as a negative regulator of inflammation, its effect on PPARγ would bring more benefits.

PPARγ functional deficiency is common in rheumatic subjects, and implicated in many RA-related pathological changes [16]. This abnormality doesn’t only occur in joints, and in fact PPARγ down-regulation is a key factor driving the imbalanced differentiation of immune cells [16-17]. These factors make PPARγ a promising anti-rheumatic target. Many compounds with PPARγ activating potentials have been confirmed with good therapeutic effects on RA, and rosiglitazone (RSG) is the most well-known representative [18]. Thanks to the pleiotropic roles of PPARγ, the related treatments can cause series of outcomes in rheumatic subjects, including angiogenesis inhibition [18-19]. Considering the crucial impacts of immune cells on RA pathology, it is apparent that the anti-angiogenesis effects of these agents are immune dependent.

Inspired by the above clues, it is reasonable to assume that QLY anti-rheumatic treatments may ease immune disruption-related angiogenesis by activating PPARγ. In this study, we further ascertained the involvement of PPARγ activation during QLY treatment on adjuvant-induced arthritis (AIA) rats by comparing its effects to RSG, a PPARγ agonist. Importantly, we identified T cells as its preferential targeting immune cells, and revealed that via this approach, potential PPARγ agonists in QLY improved the overall immune environment, and consequently inhibited synovial angiogenesis.

## 2. Materials and Methods

### 2.1. Chemicals and reagents

Incomplete Freund’s Adjuvant (IFA) and Bacillus Calmette-Guérin (BCG) were purchased from Chondex (Redmond, WA) and Rebio Scientific (Shanghai, China), respectively. Rheumatoid factor (RF), hypoxia inducible factor-1(HIF-1), and arginase-1 (ARG-1) ELISA kits were supplied by Jianglai Biotechnology (Shanghai, China). Interleukin 1 beta (IL-1β), IL-6, IL-10, IL-17A, transforming growth factor beta1 (TGF-β1), interferon gamma (IFNγ), vasoactive endothelial growth factor (VEGF), intercellular cell adhesion molecule 1(ICAM-1), C-X-C motif ligand 1 (CXCL1), monocyte chemotactic protein 1 (MCP-1), and insulin-like growth factor 1 (IGF-1) ELISA kits were supplied by Multi-Science (Hangzhou, Zhejiang, China). Glucose, lactic acid, pyruvic acid, triglyceride (TG), nonestesterified fatty acid (NEFA), reduced glutathione (GSH), malonaldehyde (MDA), inducible nitric oxide synthase (iNOS), catalase (CAT), NADH oxidase (NOX), alkaline phosphatase (AKP), and aspartate aminotransferase (AST) quantification kits were bought from Jiancheng Bioengineering Institute (Nanjing, Jiangsu, China). ReverAid first-strand cDNA synthesis kits and universal qPCR master mix were purchased from Thermo Fisher Scientific (Rockford, IL, USA) and New England Biolabs (Ipswich, MA, USA), respectively. Gene-specific primers used were synthesized by General Biology (Chuzhou, Anhui, China). Primary β-ACTIN, GAPDH, TLR4, STAT3, P-STAT3, JNK, p-JNK, CPT1A, CD36, PPARγ, and ATGL antibodies were obtained from ABclonal Technology (Wuhan, Hubei, China) or Affinity Biosciences (Changzhou, Jiangsu, China). HRP-conjugated IgG antibodies were purchased from Boster Biological Technology (Wuhan, Hubei, China). Red donkey anti-rabbit and orange donkey anti-mouse secondary antibodies were bought from Abbkine (Wuhan, Hubei, China). PE-tagged CD172a, APC-tagged CD43, PE-tagged CD16, FITC-tagged CD3, PE-tagged CD4 antibodies were procured from BioLegend (SanDiego, CA, USA) or Invitrogen (Carlsbad, CA, USA). RPMI-1640 medium, polyformaldehyde, crystal violet staining solution were procured from Biosharp (Beijing, China). Fetal bovine serum (FBS) was the product of Wisent (Montreal, Quebec, Canada). Pure compounds sinomenine, berberine, and palmatine were purchased from Bencao Yikang (Nanjing, Jiangsu, China). T0070907 (a PPARγ selective inhibitor) was purchased from Aladdin (Shanghai, China). Rat peripheral blood lymphocyte isolation kit and rat/human peripheral blood monocyte isolation kit were procured from Solarbio (Beijing, China). Fluorescent dye CFDA-SE and matrigel were bought from from MedChemExpress (Monmouth Junction, NJ, USA) and Thermo Fisher Scientific (Rockford, IL, USA), respectively.

### 2.2. AIA induction and treatments

The *in vivo* experiments were based on Wistar rats, which were supplied by Tianqin Biotechnology (Changsha, Hunan, China). They were kept in a qualified SPF lab with the standard conditions, and fed by the commercial rodent chow and boiled tap water. All invasive operations were conducted under anesthesia by the help of pentobarbital sodium (30 mg/kg). The greatest efforts were devoted to minimize the experimental animals’ suffering.

AIA induction was conducted based on a previously detailed protocol using male rats (8 weeks-old) [11-13]. IFA and BCG were mixed to prepare complete Freund’s adjuvant (CFA), which was used to provoke immune reactions in all the rats except those assigned into normal group. Since CFA injection, the immunized rats were divided randomly into 3 groups, and received various treatments for 30 days. Each group contained 5 rats, and the grouping is as below: healthy controls (normal, treated by 0.5% CMC-Na), AIA controls (AIA, treated by 0.5% CMC-Na), QLY-treated AIA rats (QLY, treated by QLY extract dispersed in 0.5% CMC-Na, equivalent to 10 g/kg of crude drugs), and RSG-treated AIA rats (RSG, treated by RSG resolved in 0.5% CMC-Na, 2 mg/kg). All the treatments were administrated daily by gavage. During the treatment period, arthritis scores and body weights were periodically recorded. The qualification criterion of arthritis scores was based on a previous report [20].

### 2.3. Therapeutic outcomes evaluation

One hour after the last administration on day 30, all the rats were anesthetized. The maximum amounts of blood were collected through abdominal aorta until the animals were euthanized. Blood plasma and serum were prepared by centrifugation. Levels of representative cytokines, metabolites and diagnostic indicators of RA in blood were detected by commercial quantification kits. Rat monocytes and lymphocytes were prepared using anti-coagulation blood by the aid of isolation kits. Blood serum and immune cells were used in the *in vitro* experiments. Main organs were dissected and weighted. Joint injuries were assessed by histological examinations with the reported protocols [21-24]. The livers were homogenized. Levels of metabolites and cytokines in the homogenates were detected by the corresponding kits. Expression of certain immune and metabolism-related proteins in the livers and spleens was studied by western-blot (WB) method with routine procedures [23-25].

### 2.4. Assessment of QLY-caused impacts on monocytes and lymphocytes

Distribution of monocyte subsets was investigated by flow cytometry (FCM) method [11, 26]. The whole blood collected above were spiked into 6-fold volumes of red cells lysis buffer, incubated, and centrifuged. The precipitates were re-suspended, and incubated with fluorescein-tagged antibodies (APC-CD43 and PE-CD172a) in dark. The stained cells were further washed by PBS, filtered (200 mesh), and fed to a BD FACSVerse instrument. Some of the isolated monocytes were cultured in complete RPMI-1640 medium for 24 h. Levels of certain cytokines in the medium were determined by ELISA kits. The remaining medium was used to incubate healthy rat lymphocytes, whose status was evaluated by polymerase chain reaction (PCR) with routine procedures [27-28]. The primed used were synthesized by Universal Biology (Chuzhou, Anhui, China), and their sequences are shown in **Supplementary S1**. Afterwards, we cultured the lymphocytes isolated from QLY-treated or AIA model rats for 24 h, and detected representative cytokines in the medium. The remaining medium was used to culture healthy rat monocytes, and their secretion capacity was evaluated by ELISA tests.

Next, we treated normal monocytes with different blood serums. To prepare QLY-containing serum, 5 healthy rats were treated by QLY for 3 consecutive days at the dose mentioned above. Five untreated rats from the same batch were killed at the same day to harvest blood serum. After normal rat monocytes were cultured in these serums for 24 h, levels of IL-1β and IL-6 in the medium were determined by ELISA kits. The rinsed cells were treated by Triton X-100, bovine serum albumin, primary antibodies (IL-1β and PPARγ), and fluorescein-tagged secondary antibodies in turns. Finally, cell nucleus was stained by DAPI [29]. In a replicate experiment, RA patients’ monocytes were used instead. In addition to ELISA analysis, CD16^+^ cell subset was analyzed by FCM method in this experiment.

### 2.5. Blood transfusion-based angiogenesis experiments

Six pairs of sex-matured male and female rats were kept separately for breeding. The pups were raised for 8 weeks. By then, 10 male offspring were induced with AIA, and half of them received QLY treatment. When inflammation became obvious in AIA rats (day 14), 1 ml of peripheral blood was sampled from each donator through fossa orbitalis vein every 3 days, and injected into caudal vein of their female littermates.

To investigate distribution of the implanted immune cells, whole blood from an AIA rat was used to prepare peripheral blood mononuclear cells based on a gradient centrifugation method, which were then dyed with CFDA-SE. Since cell injection, the rat was observed by an Optima Max-XP instrument at pre-determined intervals. The excitation and emission wavelengths were set at 500 and 520 nm, respectively.

On the day of the 8^th^ blood injection, 0.4 ml of matrigel supplemented with VEGF (10 ng/ml) was injected subcutaneously on abdomen [30]. The matrigel was thawed under 4° overnight beforehand. Three days later, the recipients were sacrificed. The formed plugs were retrieved, photographed, and subjected to H&E histological examination. Blood was collected through abdominal aorta for FCM analysis. Hinder paws were homogenized. Levels of cytokines and biochemical indicators in rats’ knee homogenates and remaining blood samples were determined by corresponding kits.

### 2.6. Evaluating immune-mediated anti-angiogenesis effects of compounds

To precipitate proteins, 100 μl of QLY-containing serum was spiked into 400 μl of methanol [31-32]. After a centrifugation (12000 rpm for 10 min), the supernatant was injected into a 1260 Agilent Infinity II LC system for quantification analysis. The separation was achieved on a Waters C18 column (4.6 × 250 mm, 8 μm). A gradient elution program was adopted, and sodium dihydrogen phosphate aqueous solution (10 mM, pH = 8.0) and acetonitrile served as mobile phases A and B respectively: 0-5 min, 20% A; 5-10 min, 50% A; 10-15 min, 20% A. The flow rate, injection volume and detection wavelength were set at 1 ml/min, 20 μl and 270 nm, respectively. The quantification curves for sinomenine, berberine and palmatine were developed by analyzing a serial of diluted solutions, which contained all the reference compounds.

THP-1 and Jurkat cells were cultured and passaged under ordinary conditions [33]. They were seeded in 6-wells plates. After 12 h adaptive growth, the supernatant was replaced by fresh RPMI-1640 medium supplemented with 10% normal healthy or AIA rat serums. Some of the AIA serum-stimulated cells were treated by a mixture (sinomenine, berberine and palmatine) at various concentrations. Twenty-four hours later, some cytokines in medium were detected, and the mediums were collected.

HUVEC cells were seeded in a 6-wells plate at an appropriate density. After the adaptive culture, they were incubated in the medium from immune cells culture. In this experiment, only one optimized concentration of the compounds mixture was adopted when treating immune cells, and one of the treatment wells was added by T0070907 (18 μM). HUVEC cells monolayer was scratched by a pipette tip. Twelve hours later, the wound was observed again, and VEGF levels in the medium were detected [30]. Next, we seeded HUVEC cells in a matrigel-coated 6-wells plate. After attachment, they were treated by the same medium and chemical as above for 24 h. Capillary tubules were observed and photographed [30]. In a replicate experiment, HUVEC cells were seeded in matrigel-coated transwells (8 μm). The lower chambers were filled with the medium mentioned above. After 24 h, the cells infiltrated through pores were stained with crystal violet and photographed.

### 2.7. Statistical analysis

All the data were presented as mean ± standard deviation. Statistical differences among groups were analyzed based on T-test or one-way analysis of variance coupled with Tukey post hoc test using GraphPad Prism 8.0 (GraphPad Software, Cary, NC, USA).

## 3. Results

### 3.1. QLY and RSG alleviated AIA-related angiogenesis in rats

AIA-related articular manifestations reach peak about day 14. Since then, therapeutic effects of the two treatments became obvious, causing profound decrease in arthritis scores. But they didn’t prevent AIA-caused decrease in body weight gain (**Figure 1A**). By the end of the *in vivo* experiment, paw swelling was still evident in AIA rats, and this situation was greatly eased in all the treated rats (**Figure 1B**). Among the main organs, liver and spleen showed the most significant weight changes. Hepatomegaly and splenomegaly were obvious in AIA rats, which were greatly reduced by QLY and RSG treatments (**Figure 1C**). This serves as a sign of the improved immune status, which was solidly confirmed by ELISA analysis.

**Figure 1.**
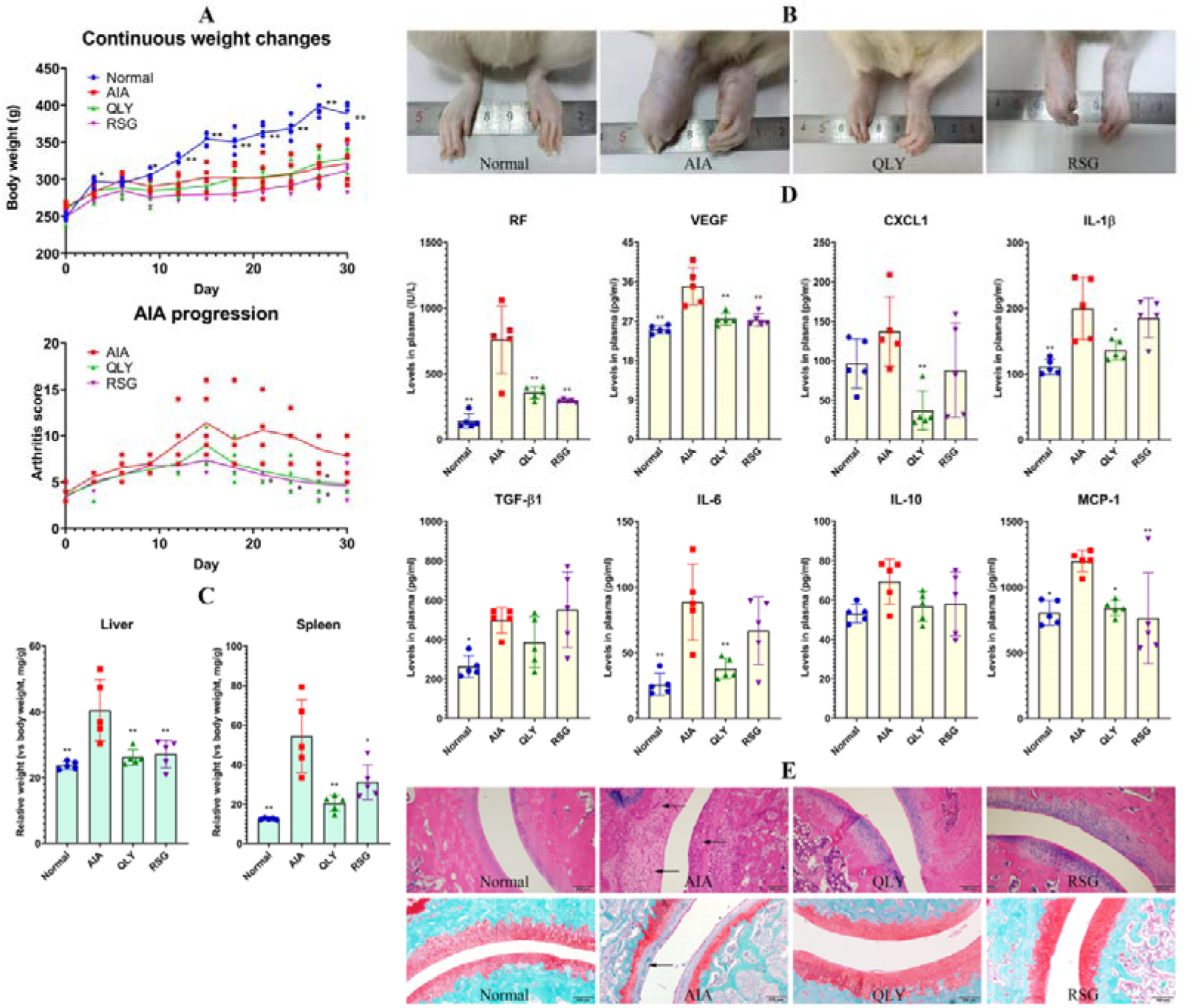
QLY reduced AIA severity in rats. **A**, periodic changes of arthritis scores and body weights; **B**, morphological changes of hind paws; **C**, relative weights of liver and spleen; **D**, levels of RA diagnostic indicators in blood; **E**, joint injuries in hind knee joints (arrows in the above H&E staining sections indicate angiogenesis; arrows in the below SOFG staining sections indicate cartilage erosion). Statistical significance: *p < 0.05 and **p < 0.01 compared with AIA model rats.

Compared to normal controls, all the tested RA-related indicators in blood were elevated in AIA rats. RF levels were increased by approximately 4-fold, indicating the successful duplication of RA models. The increase in levels of CXCL1, IL-1β, IL-6 and MCP-1 shows inflammatory activation of immune system. Interestingly, levels of IL-10 and TGF-β1, two anti-inflammatory cytokines, were also up-regulated. VEGF increase in AIA rats demonstrates an environment favorable for angiogenesis. All the abnormal changes were improved by the treatments. Although QLY performed better than RSG in controlling CXCL1, IL-1β and IL-6 levels, they were similarly effective in reducing VEGF (**Figure 1D**). Benefited from that, the two treatments both greatly reduced angiogenesis in the joints of AIA rats (H&E staining sections). As the result, joint degradation was prevented (SOFG staining sections) (**Figure 1E**). These facts above suggest that the therapeutic actions of QLY were similar to RSG, and they can both improve immune environment and ease abnormal angiogenesis in AIA rats.

### 3.2. QLY and RSG relieved AIA-related metabolic abnormalities by activating PPARγ

Because PPARγ controls metabolism, we observed metabolic changes in the treated rats to ascertain the effects of the treatments on this signaling. We observed lactic acid and pyruvic acid increase as well as glucose decrease in AIA rats, demonstrating the accelerated glycolysis, a sign of inflammation [30]. Meanwhile, TG was increased a bit. QLY and RSG generally narrowed these AIA-caused metabolic changes (**Figure 2A**). The altered metabolism conditions would impact oxidative balance *in vivo*. The increase of MDA/CAT ratio exhibits escalated oxidative stress, which was restored by QLY and RSG more or less. Confusingly, GSH levels were increased in AIA rats too, and barely affected by the two treatments (**Figure 2B**). IGF-1 is a novel RA-related indictor, controlling glucose and mineral metabolism [34]. It levels were decreased under AIA conditions. But neither of the treatments improved this situation (**Figure 2C**).

**Figure 2.**
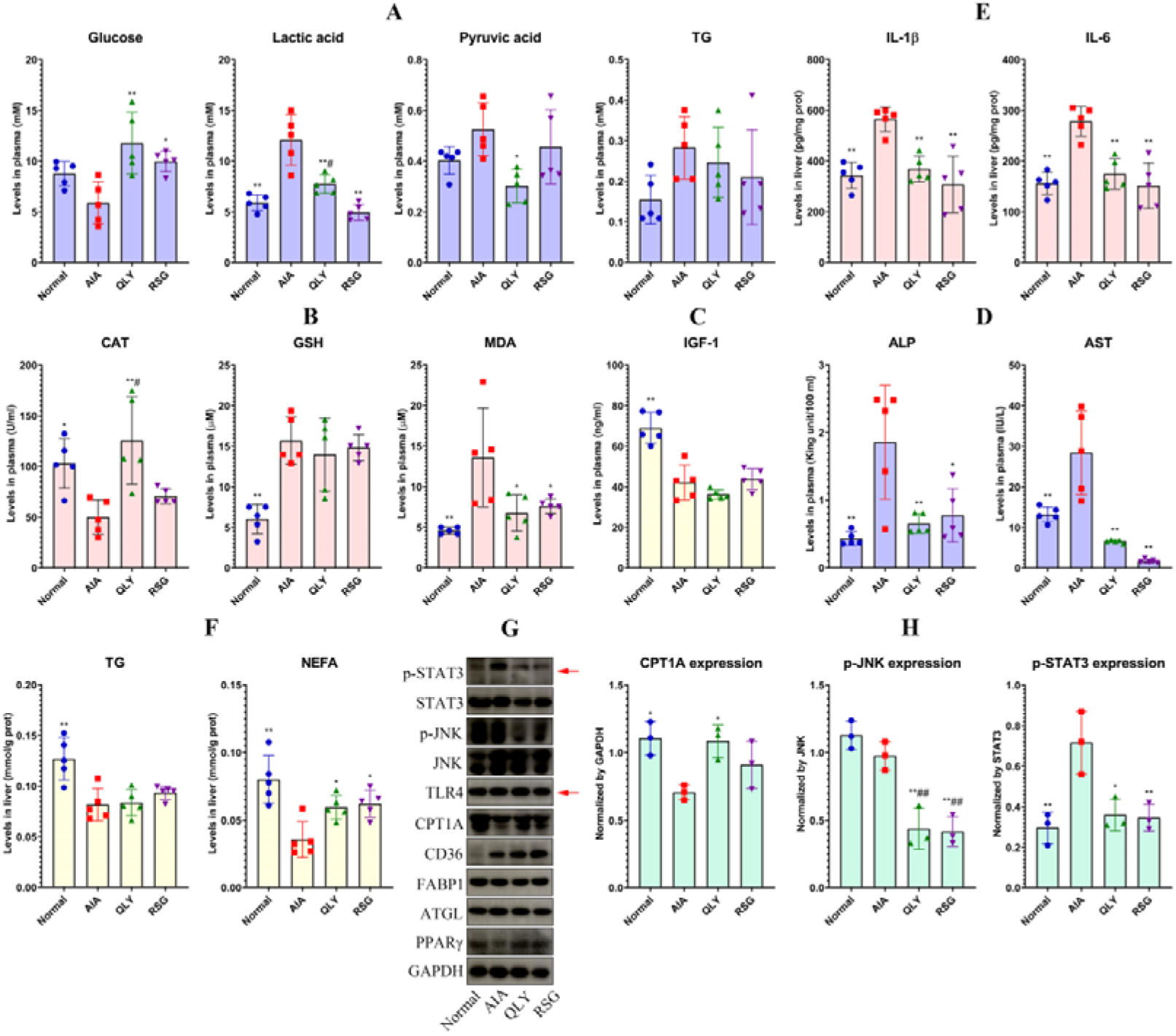
QLY improved metabolic disorders in AIA rats by activating PPARγ. **A**, levels of representative metabolites in blood; **B**, levels of oxidative indicators in blood; **C**, levels of IGF-1 in blood; **D**, levels of liver injuries indicators in blood; **E**, levels of inflammatory cytokines in liver; **F**, levels of representative lipid metabolites in liver; **G**, WB assay performed using liver samples; **H**, the quantification results of assay G. Statistical significance: *p < 0.05 and **p < 0.01 compared with AIA model rats; ^#^p < 0.05 and ^##^p < 0.01 compared with normal healthy rats.

Because metabolism is largely controlled by the liver, we next focused changes there. ALP and AST levels were increased a lot, which serve the solid evidence for hepatic injury (**Figure 2D**). It was attributed to AIA-related systemic inflammation, and levels of IL-1β and IL-6 were increased significantly in AIA rats’ liver (**Figure 2E**). After QLY and RSG treatments, inflammatory liver injuries were thoroughly cured (**Figure 2D-E**). These changes must affect metabolism functions of the liver. Compared to the normal controls, TG and NEFA levels were decreased a lot in the liver of AIA rats. Although the treatments didn’t affect TG, they effectively restored NEFA levels (**Figure 2F**). To clarify the reason accounting for the metabolic changes there, we performed WB analysis (**Figure 2G**). All the uncropped raw band images were included in **Supplementary S2**. We observed that expression of ATGL, CD36, and CPT1A was unchanged, up-regulated and down-regulated in AIA rats’ liver, respectively (**Figure 2H**). It can be inferred that liver took in more NEFA, but utilized less under AIA conditions. Meanwhile, fat mobilization remained stable. Hence, the reduced TG and NEFA must be caused by the impaired synthesis, which reflects substantial deficiency of PPARγ functions. Indeed, compared to the normal controls, PPARγ expression in AIA rats’ liver was reduced. Similar to RSG, QLY restored its levels, and promoted CPT1A expression, a downstream target of PPARγ (**Figure 2H**). This experiment also provided some useful clues to explain the reason for AIA-related hepatic inflammation. TLR4 as well as its downstream JNK were barely affected by AIA conditions, while STAT3 phosphorylation was greatly promoted (**Figure 2G**). Hence, Th17 cells but not monocytes/macrophages may initiate inflammation there [35]. RSG and QLY greatly reduced p-STAT3 expression, but showed weak effects on TLR4/JNK pathway. It suggests that PPARγ regulation may exert more impacts on T cells than monocytes/macrophages.

### 3.3. T cells were preferential targets of QLY treatment in AIA rats

In addition to T cells, monocytes/macrophages also play a crucial role in RA-related inflammatory manifestations, and are important anti-rheumatic targets [36-37]. We performed FCM analysis to investigate the impacts of QLY and RSG treatments on monocytes (**Figure 3A**). It was revealed that non-classic CD43^+^CD172a^+^ monocyte subset was increased by 3-fold in AIA rats, but classic CD43^+^CD172a^-^ monocytes were reduced. Both the treatments restored these abnormal changes (**Figure 3B**). Also, they showed potentials in down-regulating NOX and iNOS levels in the blood of AIA rats, two indicators of inflammatory monocytes/macrophages (**Figure 3C**) [37]. To confirm whether immune phenotype of these cells was fundamentally changed, we isolated them from blood and cultured *in vitro*. AIA rat monocytes were distinctly different from the normal controls. They released more IL-1β, IL-6, TGF-β1 and IL-10. However, no improvement concerning this was observed in the monocytes isolated from QLY and RSG-treated AIA rats (**Figure 3D**). This fact questioned if these PPARγ activating treatments can substantially affect monocytes/macrophages.

**Figure 3.**
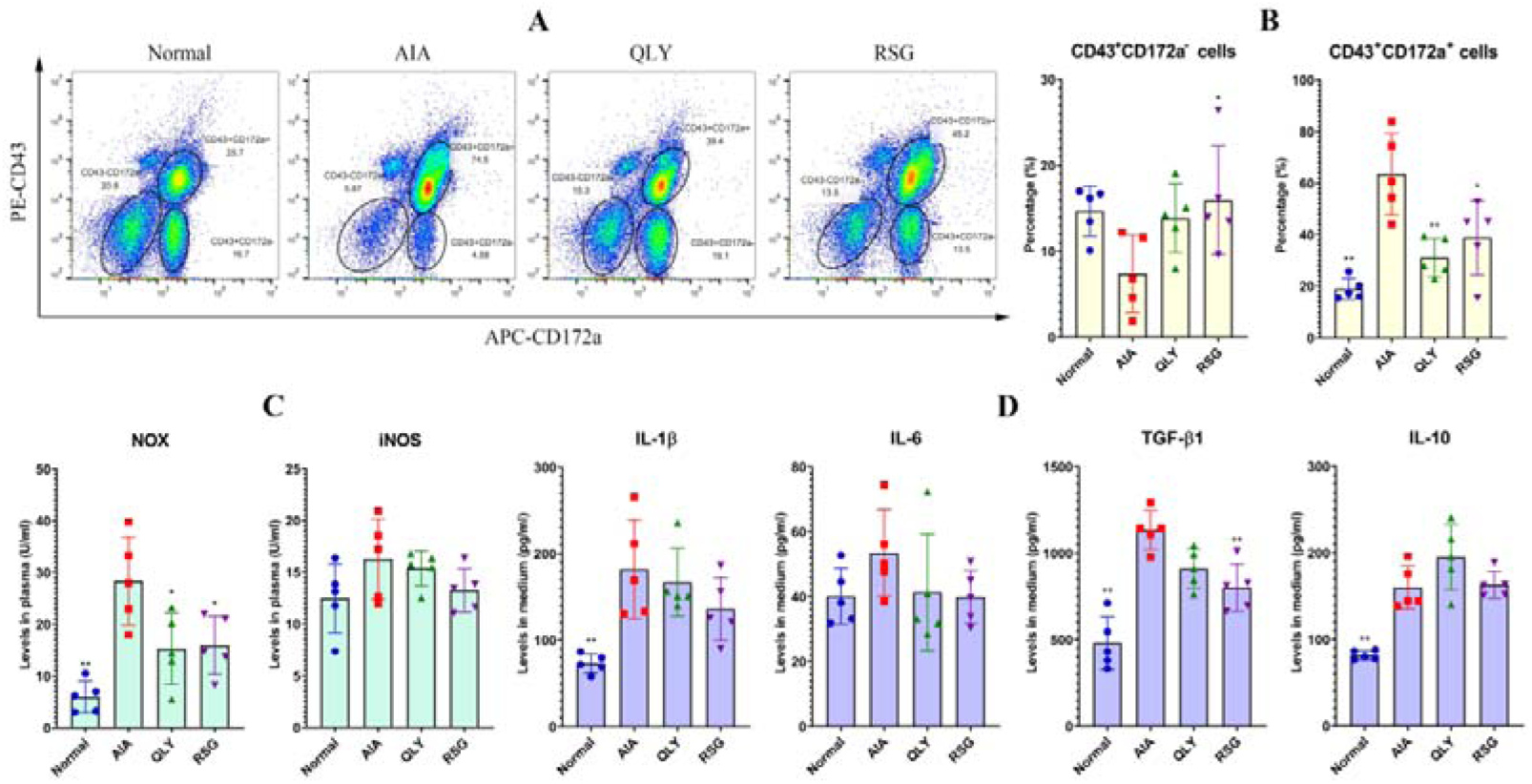
QLY-induced monocyte phenotype changes were unsustainable *in vitro*. **A**, FCM analysis of different monocyte subsets in the rats’ blood; **B**, the quantification results of assay A; **C**, levels of inflammatory monocyte indicators in blood; **D**, levels of cytokines released by the *in vitro* cultured rat monocytes. Statistical significance: *p < 0.05 and **p < 0.01 compared with AIA model rats.

Next, we co-cultured normal rat lymphocytes with the monocytes from either QLY-treated or model AIA rats, and performed PCR analysis using the lymphocytes. This experiment confirmed that QLY didn’t fundamentally change immune nature of AIA monocytes, because the monocytes from QLY-treated AIA rats showed similar impacts on expression of different Th cell subsets-related genes (*IL-6*, *IFN*γ, *IL-10*, *IL-17A*, *FOXP3*, and *ROR-*γ*T*) to those from AIA models (**Figure 4A**). In accordance to this conclusion, we found QLY-containing rat serum showed no difference with the normal serum on IL-1β and IL-10 secretion of AIA monocytes *in vitro* (**Figure 4B**). Immunofluorescence assay obtained a similar result. QLY-containing serum neither reduced IL-1β secretion nor promoted PPARγ expression (**Figure 4C**). We noticed that the expression of IL-1β and PPARγ was not always relevant. Some cells could highly express IL-1β and PPARγ at the same time. It further confirms that monocytes are not preferential targets of PPARγ agonists. To confirm this, we repeated the above cell culture by using RA patients’ monocytes. Once again, QLY-containing rat serum showed no effect on the immune indicators IL-1β, IL-10, IL-6 and ARG-1 (**Figure 4D**). FCM analysis shows than it even increased the distribution of CD16^+^ subset in the cultured human monocytes, the traditionally defined inflammatory cells (**Figure 4E-F**) [38]. Regardless of this mystery, we can conclude that QLY treatment exert no anti-inflammatory effects on monocytes directly at treatment doses.

**Figure 4.**
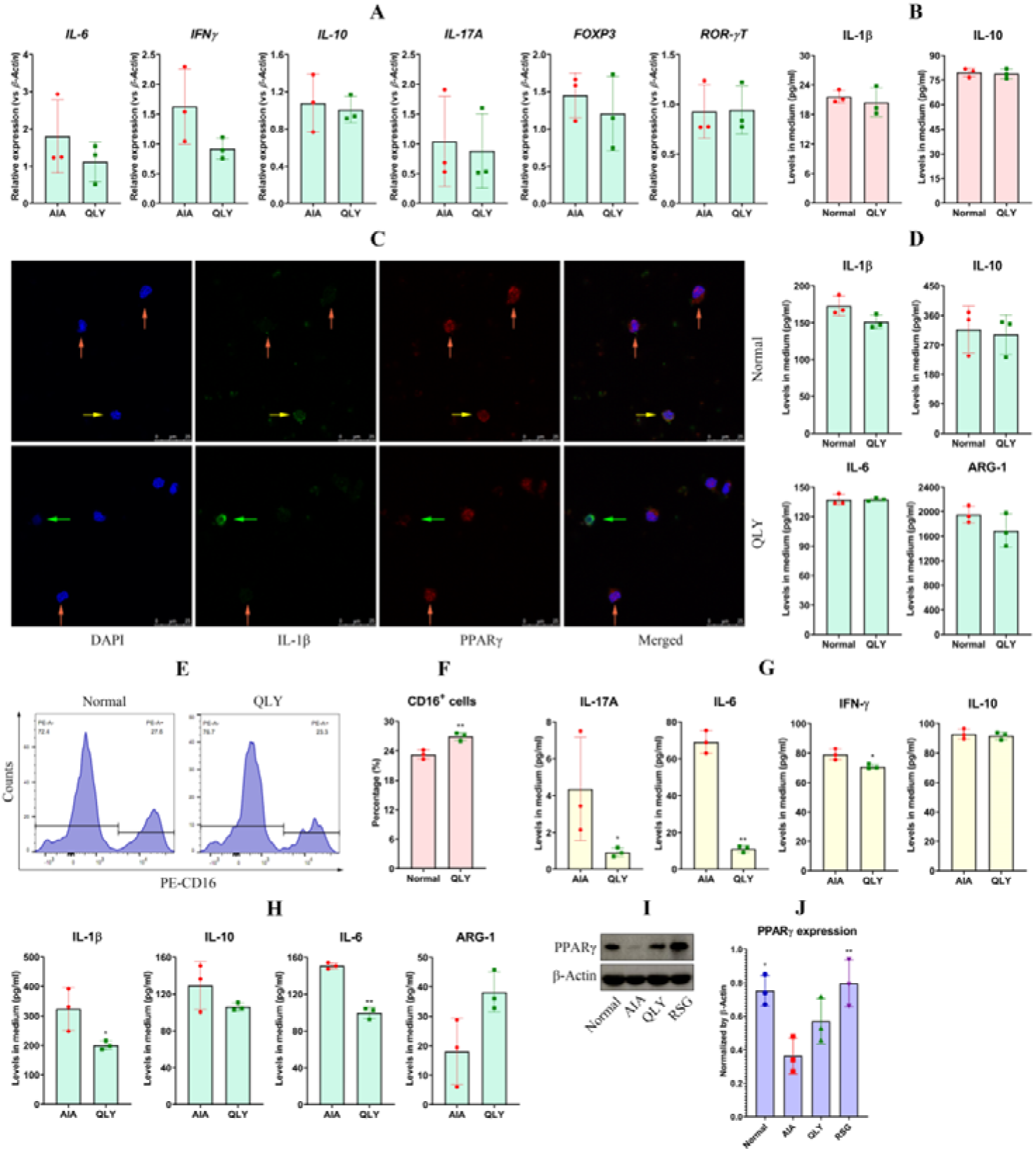
QLY preferentially targeted inflammatory T cells when treating AIA rats. **A**, expression of various Th subsets-related genes in the normal lymphocytes grew in the monocyte culture medium from the experiment above; **B**, levels of cytokines released by AIA rats’ monocytes cultured by normal or QLY-containing serum; **C**, expression and co-localization of IL-1β and PPARγ in these monocytes; **D**, levels of cytokines released by RA patients’ monocytes cultured by normal or QLY-containing serum; **E**, FCM analysis of these cells; **F**, the quantification results of assay E; **G**, levels of cytokines released by the *in vitro* cultured lymphocytes from different rats; **H**, levels of cytokines released by normal monocytes grew in the lymphocyte culture medium above; **I**, WB assay performed using spleen samples; **J**, the quantification results of assay I. Statistical significance: *p < 0.05 and **p < 0.01 compared with AIA group.

Hence, the observed changes of monocytes in treated AIA rats must be mediated by T cells, the dominate role in RA pathology [2]. As anticipated, the lymphocytes isolated from QLY-treated AIA rats released much less IL-17A, IL-6 and IFN-γ than the model controls, indicating impaired Th1/17 differentiation. IL-10 levels remained unchanged, showing unaffected Treg differentiation (**Figure 4G**). Furthermore, it was revealed that compared to the normal monocytes co-cultured by AIA rat lymphocytes, those co-cultured by the lymphocytes from QLY-treated AIA rats produced less IL-1β, IL-10, and IL-6. The opposite outcome occurred on ARG-1 levels (**Figure 4H**). These evidences confirm that QLY inhibited inflammatory polarization of monocytes by inhibiting Th1/17 differentiation. Spleen is the immune organ where lymphocytes reside and mature. We performed a WB assay using these samples (**Figure 4I**), and observed that PPARγ expression in AIA rats’ spleen was reduced dramatically, which was restored by QLY and RSG to certain extents (**Figure 4J**). It validates the claim that QLY reshapes rheumatic subjects’ T cells phenotypes by activating PPARγ, and improves overall immune milieu by affecting other immune cells differentiation.

### 3.4. T cells-mediated anti-angiogenesis effects of QLY in AIA rats

To test the central role of T cells in QLY anti-rheumatic treatment, we transplanted the peripheral immune cells from QLY-treated or AIA model rats in healthy recipients and compared the consequences. During the intermittent whole blood transfusion, the impacts from short-lived granulocytes would be negligible. Because of much larger population size, lymphocytes will bring more obvious impacts than monocytes on the recipients. The immune cells from AIA rats’ blood were first widely distributed in the body through circulation channels, and viscera, spine and joints were their enrichment sites. Gradually, the cells in chest were eliminated. The remaining cells were enriched in knee joints and spine (**Figure 5A**). It exhibits that T cells were the main long-lived immune cells from AIA rats’ blood functional in the recipients, became they are autoimmune reactive to cartilage [2]. Similar to the outcomes above, QLY-treated AIA rats’ blood transfusion caused a decline in CD43^+^CD172a^+^ cell counts in the recipients, albeit the change wasn’t significant (**Figure 5B-C**). We analyzed a panel of cytokines to better demonstrate immune cells transplant-caused consequences. Disappointingly, blood transfusion didn’t cause notable systematic changes. In fact, the counts of CD3^+^CD4^+^ T cells in the recipients of the two groups were also similar (**Supplementary S3**). Among all the investigated indicators, only IL-1β and ARG-1 were found changed in the recipients receiving QLY-treated AIA rats’ blood (**Figure 5D**). The decrease in IL-1β and increase in ARG-1 exhibit the rebalanced monocytes polarization [37]. But iNOS levels reduction in the recipients’ blood wasn’t obvious (**Figure 5E**). There was no difference about MDA, GSH and CAT levels between the two groups, further showing their similar immune and metabolism conditions (**Figure 5F**). These results basically show that the ability of QLY-treated AIA rats’ lymphocytes in inducing inflammatory polarization of monocytes was impaired a bit.

**Figure 5.**
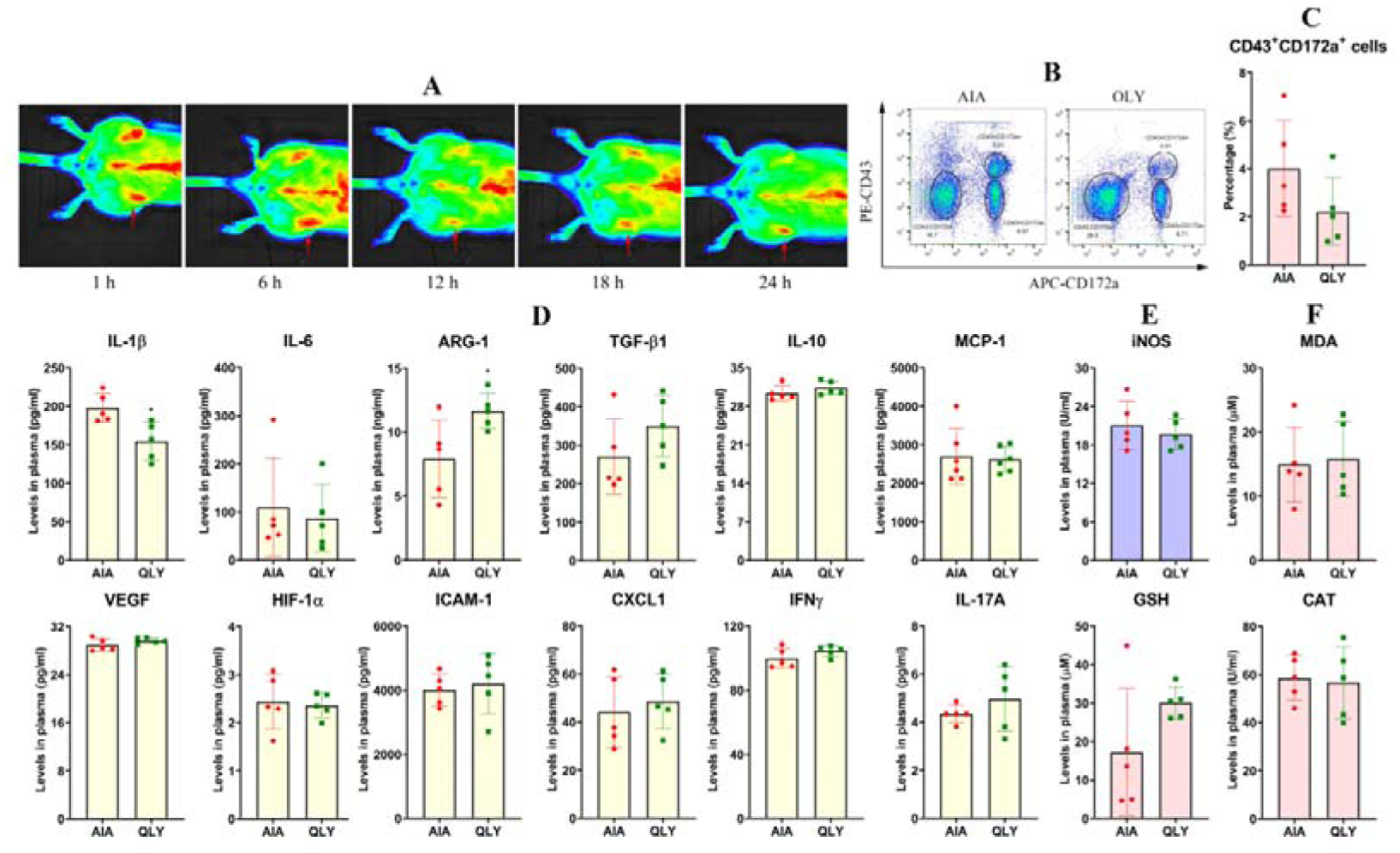
Transfusion using different rats’ blood brought little impacts on the internal environment of healthy recipients. **A**, the dynamic distribution of AIA rats’ peripheral immune cells in the recipient after transplant; **B**, FCM analysis of monocyte subsets in the recipients’ blood; **C**, the quantification results of assay B; **D**, a comprehensive evaluation of the internal environment changes by determining a panel of cytokines in blood; **E**, levels of iNOS in the recipients’ blood; **F**, levels of oxidative indicators in the recipients’ blood. Statistical significance: *p < 0.05 compared with AIA group.

Because the donators’ immune cells were enriched in joints, we believed that changes there would be more obvious. Hence, we repeated the above analyses using knee joint homogenates, which basically verified the hypothesis. Level differences of IL-1β and ARG-1 in joints between the groups were similar to those found in blood. TGF-β1 decrease didn’t only indicate altered immune conditions, but also promised the eased angiogenesis in the recipients receiving QLY-treated AIA rats’ blood [3-4]. In line with this, some other angiogenesis-related cytokines including VEGF, HIF-1α and ICAM-1 were also reduced. IL-17A and IFNγ decrease in these animals confirm that Th1/17 differentiation was hampered by QLY therapy (**Figure 6A**). Due to T cells enrichment in joints, functional difference of monocytes/macrophages between the two groups became obvious there. QLY group showed much lower levels of iNOS expression than AIA group (**Figure 6B**). Despite of these immune changes, oxidative stress was unaffected (**Figure 6C**). The above ELISA tests confirm that angiogenesis is a T cells-mediated event. To vividly demonstrate the effects brought by the immune cells transplant, we planted a matrigel plug in the healthy recipients. Obviously, AIA rats’ immune cells caused angiogenesis, indicated by dark red color of the plug, and their potentials were greatly weakened after QLY therapy (**Figure 6D**). Examination on H&E stained sections revealed dense new vessels in the plug planted in healthy recipients transfused with AIA models’ blood. A large amount of immune cells were infiltrated through these channels. These situations were improved in QLY group (**Figure 6E**). The above results show that it is plausible that QLY reshaped T cells, and then exerted anti-angiogenesis effects in AIA rats.

**Figure 6.**
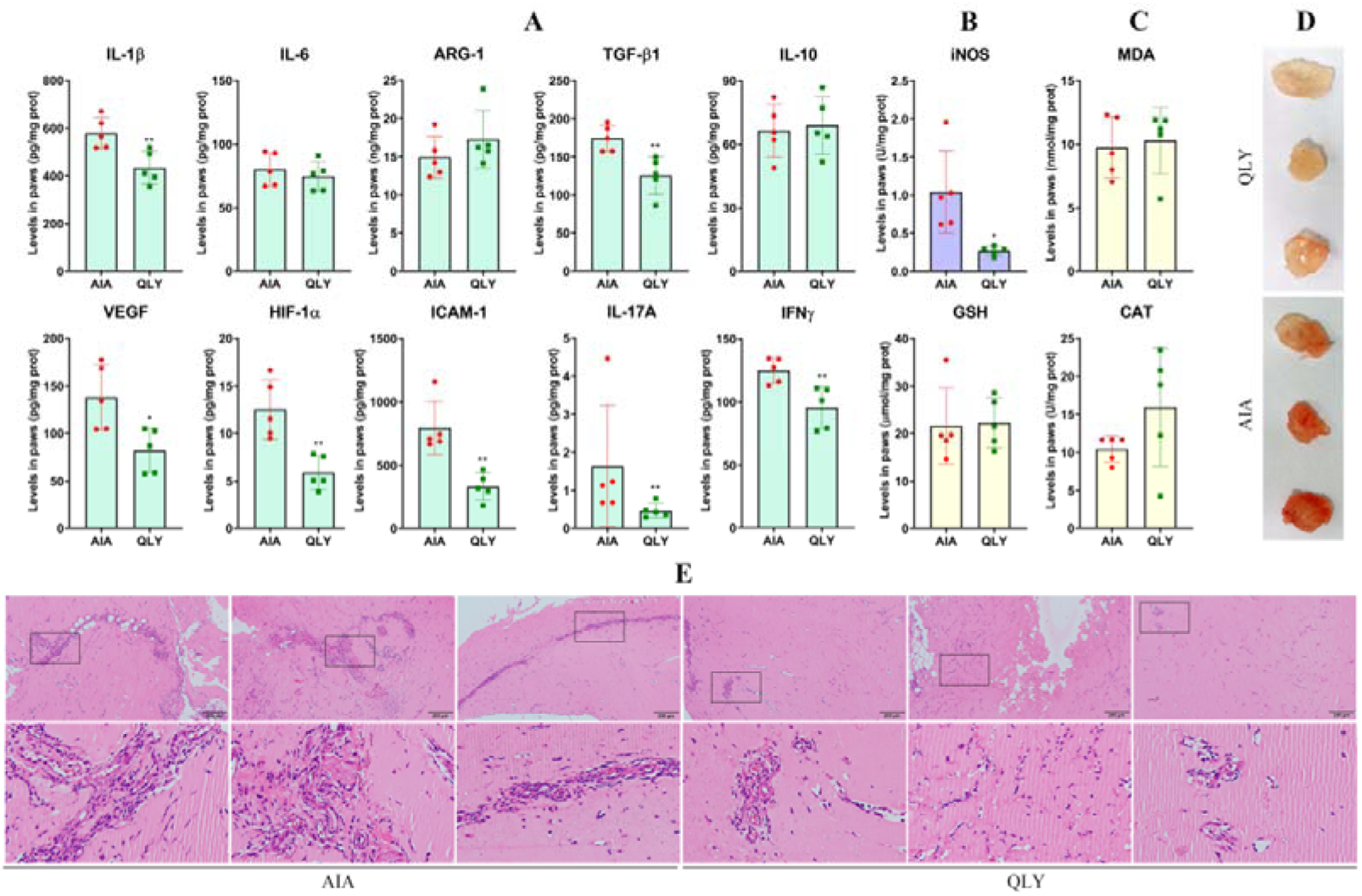
QLY impaired the ability of AIA rats’ lymphocytes inducing angiogenesis *in vivo*. **A**, levels of immune/angiogenesis-related cytokines in the recipients’ joints; **B**, levels of iNOS in the joints; **C**, levels of oxidative indicators in the joints; **D**, matrigel plugs retrieved from the recipients transfused by different rats’ blood; **E**, histological examination of H&E stained plug sections. Statistical significance: *p < 0.05 and **p < 0.01 compared with AIA group.

### 3.5. QLY-derived PPARγ agonists inhibited angiogenesis by targeting T cells

Main bioactive components in QLY are alkaloid derivatives, which contribute greatly to its anti-rheumatic effects [7-10]. In QLY-containing serum, we detected 3 alkaloids, sinomenine, berberine and palmatine (**Figure 7A**). Our unpublished study revealed that all of them are potential PPARγ agonists. Using the quantification curves (**Figure 7B**), we calculated their concentrations in the blood. The mean levels of sinomenine, berberine and palmatine were 2.08, 0.51 and 1.12 μg/ml, respectively (**Figure 7C**). The values were adopted as the medium treatment dose of the compounds mixture in following experiments. The low and high doses were divided and multiplied by 2, respectively. The mixture failed to reduce IL-1β and TGF-β1 secretion in AIA rat serum-primed THP-1 monocytes at all the three doses (**Figure 7D**). Differently, AIA serum-induced IFNγ and IL-17A production in Jurkat T cells was reduced a lot by them, generally in a dose-dependent manner (**Figure 7E**). Based on these results, we then treated rat lymphocytes at the high dose, and added a PPARγ antagonist in some treatment wells. The resulting medium was used to culture HUVEC cells. The culture medium from AIA rat serum-primed lymphocytes greatly augmented VEGF secretion in normal HUVEC cells. QLY-related compounds impaired this capacity of these rat lymphocytes, while T0070907 completely abrogated this effect (**Figure 7F**). VEGF level differences eventually affected angiogenesis potentials of HUVEC cells. Twelve hours after growth in the lymphocytes culture mediums, wound healing phenomenon was obvious in AIA group, but the compounds treatment on lymphocytes suppressed HUVEC cells migration (**Figure 7G**). Subsequently, we confirmed that the potentials of HUVEC cells infiltrating through transwell and forming tubes were reinforced when cultured by the medium from AIA rat serum-primed lymphocytes, which were inhibited by the compounds too (**Figure 7H-J**). T0070907 showed the antagonistic effects to the compounds mixture in all the above experiments. The facts confirm that T cells-mediated anti-angiogenesis effects of QLY rely on PPARγ activation.

**Figure 7.**
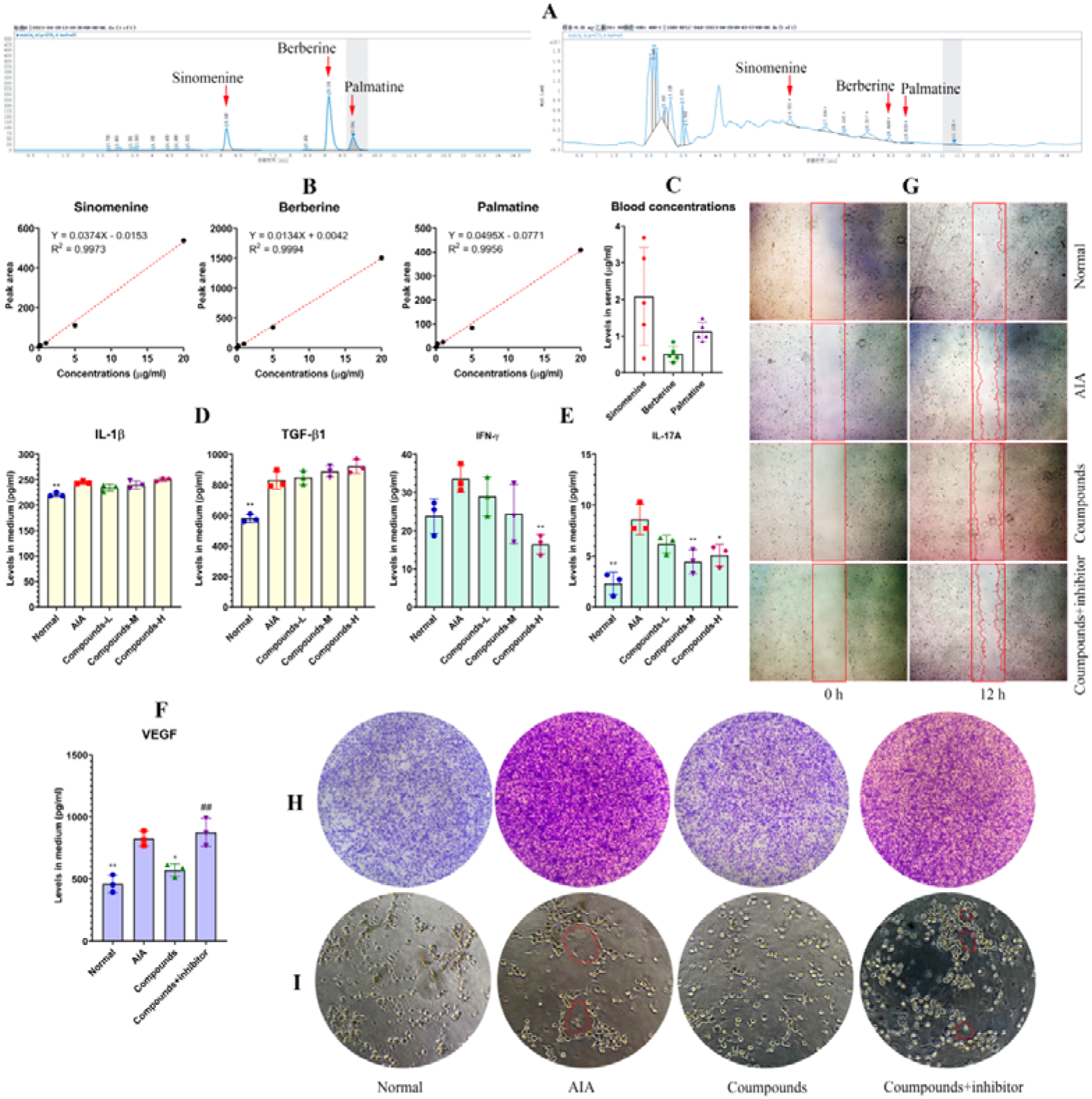
QLY-related compounds inhibited T cells-meditated angiogenesis *in vitro*. **A**, HPLC detection of the compounds in QLY-containing serum; **B**, the quantification curves developed using the reference compounds; **C**, the compounds concentrations detected in QLY-containing serum; **D**, levels of cytokines released by THP-1 cells stimulated by AIA serum and the compound mixture; **E**, levels of cytokines released by Jurkat cells stimulated by AIA serum and the compound mixture; **F**, levels of VEGF released by HUVEC cells grew in different Jurkat cells culture mediums; **G**-**H**, the impacts of different Jurkat cells culture mediums on the wound healing, transwell infiltration, and tube-forming capacities of HUVEC cells. Statistical significance: *p < 0.05 and **p < 0.01 compared with AIA group; ^##^p < 0.01 compared with compounds treatment group.

## 4. Discussion

Because of the distinctly different theories and therapeutic notions from contemporary medicine system, TCM is generally unfamiliar to the western world. Some of TCM viewpoints seem to be groundless, and lack of supporting evidence. We have to admit that being an empirical medicine, TCM always emphasize clinical efficacy rather than theory evolution. To overcome its instinct shortage, more efforts should be devoted to narrow this gap with evolving research strategy and new knowledge [39].

In RA field, humankind has entered a new era. It is clear now that immune cells are the dominant players in RA pathology, although some other cells also participate in development of RA-related manifestations. Accordingly, immune cells become the preferential targets of RA therapies. It does not only lay a foundation for developing new biological anti-rheumatic agents, but also successfully explains the therapeutic mechanisms of some conventional DMARDs. Effective therapies always achieve comprehensive improvements, but their direct targets were largely unknown. It will impose negative impacts on rational and precise use of the relevant drugs. By taking methotrexate as an example, it can ease many aspects of RA, including inflammation, angiogenesis, joints degradation, and even metabolic complications [40-41]. We can’t attribute all of the outcomes to its medicinal properties, and it is inappropriate to use this drug to treat metabolic diseases. Actually, adenosine receptors activation-caused immune changes serve as the foundation for the other effects. It is highly possible that certain natural drugs also achieve comprehensive anti-rheumatic effects by regulating immune status. Many researches support this theory [42-43].

QLY is a TCM formula showing multiple functions in treating RA [44]. The relevant researches started from investigating its effects on synovial angiogenesis [14]. But in fact, it can also treat systemic inflammation, joint injuries, and metabolic alterations. Inspired by the aforementioned clues, we believe that immune-regulating property is the key for QLY to achieve the overall anti-rheumatic outcomes. An important clue is that all the herbal components in it (Radix Sophorae flavescentis, Caulis Sinomenium acutum, Cortex Phellodendri chinensis, and Rhizoma Dioscoreae hypoglaucae) are discovered with the relevant functions [45-48].

It is a common sense that T cells drive the development and progression of RA. Meanwhile, monocytes/macrophages are the main source of inflammatory mediators and deeply implicated in RA-related inflammation. Therefore, the two representatives of adaptive and innate immune components are usually emphasized in RA researches. Our works confirmed that QLY can inhibit them from acquiring the inflammatory phenotypes [11-12]. As immune environment is shaped based on sophisticated interplay of different cells, these findings brought another puzzle: which kind of immune cells is the main targets of QLY. The factors mediating development and differentiation of T cells and monocytes/macrophages are largely different. It is not very possible that QLY is equally effective in regulating them.

One of our previous studies showed that QLY therapy hampered inflammatory differentiation of T cells and monocytes/macrophages simultaneously in rheumatic rodents [11]. Importantly, this work found that T cells were much more sensitive to QLY treatments than monocytes/macrophages at treatment doses, demonstrating the central role of T cells in QLY-induced immune changes for the first time. QLY didn’t only affect Th1/17 differentiation, but also impaired generation and maturation of T cells. The current study confirms this conclusion and provides more convincing evidence. Although QLY globally decreased inflammatory cytokines in AIA rats (**Figure 1D**), it selectively targeted T cells rather than innate immunocytes in the liver, indicated by the varied effects on STAT3 and TLR4/JNK signals (**Figure 2G**). At the same time, QLY-induced monocytes/macrophages changes cannot be sustained *in vitro* (**Figure 3D**). It hints that these outcomes relied on internal environment changes. In another word, impacts of QLY on them were indirect and mediated by some other factors. Indeed, the monocytes from AIA rats and RA patients were both unaffected by QLY-containing serum (**Figure 4**). QLY-related compounds cannot impact them too at therapeutic concentrations (**Figure 7D**). Both the *in vitro* (**Figure 4**) and *in vivo* (**Figure 5**) evidences show that T cells responded well to QLY therapy, and mediated the global immune changes. In fact as early as 2018, we had noticed that QLY would preferentially target inflammatory T cells, as it inhibited pentose phosphate pathway [10]. Since T cells initiate RA-related autoimmune reactions and most of pathological changes, this property explains many anti-rheumatic functions of QLY, including its effects on synovial angiogenisis. Meanwhile, this previous work revealed PPARγ as a key molecular target of QLY.

PPARγ is a well-known governor of lipid metabolism. Therefore, QLY-induced PPARγ activation must affect adipose tissues and liver. Previously, we showed that QLY can attenuate monocytes/macrophages-mediated inflammation in AIA rats by activating PPARγ in adipose tissues [12]. Here, we further confirm its activation effects on PPARγ by examining the changes in liver. As expected, QLY didn’t only restore the altered lipids levels there, but also directly promoted expression of PPARγ and its downstream CPT1A (**Figure 2G**). A recent study suggests that PPARγ activation during QLY treatments would be resulted from the inhibition on SIRT1 [10]. But it is difficult to explain QLY-induced PPARγ up-regulation in some cases when SIRT1 signal was normal [12]. Possibly, there exist PPARγ agonists in QLY. Our unpublished work confirmed this, and revealed that many related alkaloids in QLY are PPARγ agonist candidates. As their representatives, sinomenine, berberine and palmatine can potently inhibit inflammatory secretion of T cells, and compromised their ability in inducing angiogenesis (**Figure 7**). This finding suggests that PPARγ activation in T cells is especially beneficial in the context of RA therapies, despite of the fact that monocytes/macrophages are usually defined as PPARγ-relevant immune cells, since the transcriptional factors driving their inflammatory polarization like AP-1 and NF-κB are negatively regulated by PPARγ [16-17]. Although many studies confirm that PPARγ activation would inhibit inflammatory T cells differentiation, the detailed mechanism is still not thoroughly elucidated [49-50]. Hence, more works should be done in this aspect, which is crucial for exploiting the potentials of PPARγ-targeting anti-rheumatic therapies.

## 5. Conclusion

Similar to RSG, QLY also effectively reduced AIA severity in rats, reducing systemic inflammation, joint injuries as well as synovial angiogenesis by activating PPARγ. T cells were more sensitive to than monocytes QLY treatment. Through cell interactions, QLY improved overall immune environment by inhibiting Th1/17 differentiation. The inflammatory T cells were unable to induce obvious angiogenesis both *in vivo* and *in vitro*. The collective evidences show that QLY inhibited immune-mediated abnormal angiogenesis in AIA rats by activating PPARγ in T cells.

## Abbreviations

WAT: white adipose tissues
AIA: adjuvant-induced arthritis
QLY: Qing-Luo-Yin
RA: rheumatoid arthritis
TCM: Traditional Chinese Medicine
IFA: Incomplete Freud’s adjuvant
BCG: Bacillus Calmette-Guérin
RSG: rosiglitazone
RF: rheumatoid factor
HIF-1: hypoxia inducible factor 1
ARG-1: arginase 1
IL-1β: interleukin 1 beta
TGF-β1: transforming growth factor beta1
IFNγ: interferon gamma
VEGF: vasoactive endothelial growth factor
ICAM-1: intercellular cell adhesion molecule 1
CXCL1: C-X-C motif ligand 1
MCP-1: monocyte chemotactic protein 1
IGF-1: insulin-like growth factor 1
TG: triglyceride
NEFA: nonestesterified fatty acid
GSH: reduced glutathione
MDA: malonaldehyde
iNOS: inducible nitric oxide synthase
CAT: catalase
NOX: NADH oxidase
AKP: alkaline phosphatase
AST: aspartate aminotransferase
FBS: fetal bovine serum
PCR: polymerase chain reaction
WB: western-blot
FCM: flow cytometry
DMARD: disease modifying antirheumatic drug

## CRediT authorship contribution statement

**Jian Zuo** and **Yan Li** conceived the idea; **Meng-Ke Song** and **Qin Yin** performed majority of the experiments; **Meng-Fan Gu** and **Wen-Gang Chen** assisted all the experiments; **Jian Zuo** wrote the manuscript; **Opeyemi Joshua Olatunji** proof read the manuscript. All the authors gave final approval of the version to be published.

## Data Availability

All data generated or analyzed during this study are included in this article and its supplementary information file.

## References

1. Fukae J, Tanimura K, Atsumi T, Koike T. Sonographic synovial vascularity of synovitis in rheumatoid arthritis. Rheumatology 2014; 53(4): 586–91.

2. McInnes IB, Schett G. The pathogenesis of rheumatoid arthritis. N Engl J Med 2011; 365(23):2205–19.

3. Elshabrawy HA, Chen Z, Volin MV, Ravella S, Virupannavar S, Shahrara S. The pathogenic role of angiogenesis in rheumatoid arthritis. Angiogenesis 2015; 18(4): 433–48.

4. Wang Y, Wu H, Deng R. Angiogenesis as a potential treatment strategy for rheumatoid arthritis. Eur J Pharmacol 2021; 910: 174500.

5. Tsaltskan V, Firestein GS. Targeting fibroblast-like synoviocytes in rheumatoid arthritis. Curr Opin Pharmacol 2022; 67: 102304.

6. Smolen JS, Landewé RBM, Bergstra SA, Kerschbaumer A, Sepriano A, Aletaha D, et al. EULAR recommendations for the management of rheumatoid arthritis with synthetic and biological disease-modifying antirheumatic drugs: 2022 update. Ann Rheum Dis. 2023; 82(1): 3–18.

7. Li S, Wang R, Zhang Y, Zhang X, Layon AJ, Li Y, Chen M. Symptom combinations associated with outcome and therapeutic effects in a cohort of cases with SARS. Am J Chin Med. 2006;34(6):937–47.

8. □□□,□□. □□□□□□□□□□□□□□□□□□ [J]. □□□□□□□□□□□, 2015, 29 (06): 883–892.

9. Wang ZY, Wang X, Zhang DY, Hu YJ, Li S. [Traditional Chinese medicine network pharmacology: development in new era under guidance of network pharmacology evaluation method guidance]. Zhongguo Zhong Yao Za Zhi. 2022 Jan;47(1):7–17.

10. Zuo J, Wang X, Liu Y, Ye J, Liu Q, Li Y, Li S. Integrating network pharmacology and metabolomics study on anti-rheumatic mechanisms and antagonistic effects against methotrexate-induced toxicity of Qing-Luo-Yin. Front Pharmacol 2018; 9: 1472.

11. Wang DD, Wu XY, Dong JY, Cheng XP, Gu SF, Olatunji OJ, Li Y, Zuo J. Qing-Luo-Yin alleviated experimental arthritis in rats by disrupting immune feedback between inflammatory T cells and monocytes: key evidences from its effects on immune cell phenotypes. J Inflamm Res 2021; 14: 7467–86.

12. Wang R, Li DF, Hu YF, Liao Q, Jiang TT, Olatunji OJ, Yang K, Zuo J. Qing-Luo-Yin alleviated monocytes/macrophages-mediated inflammation in rats with adjuvant-induced arthritis by disrupting their interaction with (pre)-adipocytes through PPAR-γ signaling. Drug Des Devel Ther 2021; 15: 3105–3118.

13. Ye P, Wang QH, Liu CS, Li GH, Olatunji OJ, Lin JT, Zuo J. SIRT1 inhibitors within Qing-Luo-Yin alleviated white adipose tissues-mediated inflammation in antigen-induced arthritis mice. Phytomedicine 2024; 122: 155132.

14. Li S, Lu AP, Wang YY, Li YD. Suppressive effects of a Chinese herbal medicine Qing-Luo-Yin extract on the angiogenesis of collagen-induced arthritis in rats. Am J Chin Med 2003; 31(5): 713–20.

15. Wu YJ, Fang WJ, Pan S, Zhang SS, Li DF, Wang ZF, Chen WG, Yin Q, Zuo J. Regulation of Sirt1 on energy metabolism and immune response in rheumatoid arthritis. Int Immunopharmacol 2021; 101(Pt A): 108175.

16. Liu Y, Wang J, Luo S, Zhan Y, Lu Q. The roles of PPARγ and its agonists in autoimmune diseases: A comprehensive review. J Autoimmun 2020; 113: 102510.

17. Nobs SP, Kopf M. PPAR-γ in innate and adaptive lung immunity. J Leukoc Biol 2018; 104(4): 737–41.

18. Kwon EJ, Park EJ, Choi S, Kim SR, Cho M, Kim J. PPARγ agonist rosiglitazone inhibits migration and invasion by downregulating Cyr61 in rheumatoid arthritis fibroblast-like synoviocytes. Int J Rheum Dis 2017; 20(10): 1499–509.

19. Wagner N, Wagner KD. PPARs and angiogenesis-implications in pathology. Int J Mol Sci 2020; 21(16): 5723.

20. Zhou Y, Xiang R, Qin G, Ji B, Yang S, Wang G, Han J. Xanthones from Securidaca inappendiculata Hassk. attenuate collagen-induced arthritis in rats by inhibiting the nicotinamide phosphoribosyltransferase/glycolysis pathway and macrophage polarization. Int Immunopharmacol 2022; 111: 109137.

21. Liu JQ, Zhao XT, Qin FY, Zhou JW, Ding F, Zhou G, et al. Isoliquiritigenin mitigates oxidative damage after subarachnoid hemorrhage in vivo and in vitro by regulating Nrf2-dependent signaling pathway via targeting of SIRT1. Phytomedicine 2022; 105: 154262.

22. Zhang S, Zhu P, Yuan J, Cheng K, Xu Q, Chen W, Pan Z, Zheng Y. Non-alcoholic fatty liver disease combined with rheumatoid arthritis exacerbates liver fibrosis by stimulating co-localization of PTRF and TLR4 in rats. Front Pharmacol 2023; 14: 1149665.

23. Sun Y, Bai YP, Wang DG, Xing YJ, Zhang T, Wang W, et al. Protective effects of metformin on pancreatic β-cell ferroptosis in type 2 diabetes in vivo. Biomed Pharmacother 2023; 168: 115835.

24. Hua Z, Hui LI, Haihua W. Potential protective effects of the water-soluble Chinese propolis on experimental ulcerative colitis. J Tradit Chin Med 2023; 43(5): 925–33.

25. Wang X, Shen C, Wang X, Tang J, Wu Z, Huang Y, et al. Schisandrin protects against ulcerative colitis by inhibiting the SGK1/NLRP3 signaling pathway and reshaping gut microbiota in mice. Chin Med 2023; 18(1): 112.

26. Wu YJ, Zhang SS, Yin Q, Lei M, Wang QH, Chen WG, et al. α-Mangostin inhibited M1 polarization of macrophages/monocytes in antigen-induced arthritis mice by up-regulating silent information regulator 1 and peroxisome proliferators-activated receptor γ simultaneously. Drug Des Devel Ther 2023; 17: 563–77.

27. Yin M, Ding X, Yin S, Wang L, Zhang K, Chen Y, et al. Exosomes from hepatitis B virus-infected hepatocytes activate hepatic stellate cells and aggravate liver fibrosis through the miR-506-3p/Nur77 pathway. J Biochem Mol Toxicol 2023; 37(10): e23432.

28. Xu S, He L, Ding K, Zhang L, Xu X, Wang S, Qian X. Tanshinone IIA ameliorates streptozotocin-induced diabetic nephropathy, partly by attenuating PERK pathway-induced fibrosis. Drug Des Devel Ther 2020; 14:5773–82.

29. Zhou HH, Zhang YM, Zhang SP, Xu QX, Tian YQ, Li P, et al. Suppression of PTRF alleviates post-infectious irritable bowel syndrome via downregulation of the TLR4 pathway in rats. Front Pharmacol 2021; 12: 724410.

30. Jiang TT, Ji CF, Cheng XP, Gu SF, Wang R, Li Y, Zuo J, Han J. α-Mangostin alleviated HIF-1α-mediated angiogenesis in rats with adjuvant-induced arthritis by suppressing aerobic glycolysis. Front Pharmacol 2021; 12: 785586.

31. Liu Y, Sun H, Li C, Pu Z, Wu Z, Xu M, et al. Comparative HPLC-MS/MS-based pharmacokinetic studies of multiple diterpenoid alkaloids following the administration of Zhenwu Tang and Radix Aconiti Lateralis Praeparata extracts to rats. Xenobiotica 2021; 51(3): 345–54.

32. Li H, Wang Y, Fan R, Lv H, Sun H, Xie H, et al. The effects of ferulic acid on the pharmacokinetics of warfarin in rats after biliary drainage. Drug Des Devel Ther 2016; 10: 2173–80.

33. Cheng L, Wang H, Wang Z, Huang H, Zhuo D, Lin J. Leflunomide inhibits proliferation and induces apoptosis via suppressing autophagy and PI3K/Akt signaling pathway in human bladder cancer cells. Drug Des Devel Ther 2020; 14: 1897–908.

34. Lee H, Suh YS, Lee SI, Cheon YH, Kim M, Noh HS, Kim HO. Serum IGF-1 in patients with rheumatoid arthritis: correlation with disease activity. BMC Res Notes 2022; 15(1): 128.

35. Chen JY, Tian XY, Wei SS, Xu W, Pan RR, Chen LL, et al. Magnolol as STAT3 inhibitor for treating multiple sclerosis by restricting Th17 cells. Phytomedicine 2023; 117: 154917.

36. Yin Q, Wu YJ, Pan S, Wang DD, Tao MQ, Pei WY, Zuo J. Activation of cholinergic anti-inflammatory pathway in peripheral immune cells involved in therapeutic actions of α-mangostin on collagen-induced arthritis in rats. Drug Des Devel Ther 2020; 14: 1983–93.

37. Lei M, Tao MQ, Wu YJ, Xu L, Yang Z, Li Y, et al. Metabolic enzyme triosephosphate isomerase 1 and nicotinamide phosphoribosyltransferase, two independent inflammatory indicators in rheumatoid arthritis: evidences from collagen-induced arthritis and clinical samples. Front Immunol 2022; 12: 795626.

38. Kapellos TS, Bonaguro L, Gemünd I, Reusch N, Saglam A, Hinkley ER, Schultze JL. Human monocyte subsets and phenotypes in major chronic inflammatory diseases. Front Immunol 2019; 10: 2035.

39. Xu H, Li S, Liu J, Cheng J, Kang L, Li W, et al. Bioactive compounds from Huashi Baidu decoction possess both antiviral and anti-inflammatory effects against COVID-19. Proceedings of the National Academy of Sciences 2023;120(18): e2301775120.

40. Friedman B, Cronstein B. Methotrexate mechanism in treatment of rheumatoid arthritis. Joint Bone Spine 2019; 86(3): 301–7.

41. Zhao Z, Hua Z, Luo X, Li Y, Yu L, Li M, et al. Application and pharmacological mechanism of methotrexate in rheumatoid arthritis. Biomed Pharmacother 2022; 150: 113074.

42. Chen WG, Zhang SS, Pan S, Wang ZF, Xu JY, Sheng XH, et al. α-Mangostin treats early-stage adjuvant-induced arthritis of rat by regulating the CAP-SIRT1 pathway in macrophages. Drug Des Devel Ther 2022; 16: 509–20.

43. Wang Z, Cheng G, Wu M. Columbianetin improve rheumatoid arthritis and relative mechanisms. Lat Am J Pharm 2021; 40(8): 1910–7.

44. Zhou W, Yang K, Zeng J, Lai X, Wang X, Ji C, Li Y, Zhang P, Li S. FordNet: Recommending traditional Chinese medicine formula via deep neural network integrating phenotype and molecule. Pharmacol Res. 2021 Nov;173:105752.

45. Zhou BH, Yang JY, Ding HY, Chen QP, Tian EJ, Wang HW. Anticoccidial effect of toltrazuril and Radix Sophorae Flavescentis combination: Reduced inflammation and promoted mucosal immunity. Vet Parasitol 2021; 296: 109477.

46. Zhao XX, Peng C, Zhang H, Qin LP. Sinomenium acutum: a review of chemistry, pharmacology, pharmacokinetics, and clinical use. Pharm Biol 2012; 50(8): 1053–61.

47. Yin MC, Chang CH, Su CH, Yu B, Hsu YM. Pteris multifida, Cortex phellodendri, and probiotics attenuated inflammatory status and immunity in mice with a Salmonella enterica serovar Typhimurium infection. Biosci Biotechnol Biochem 2018; 82(5): 836–47.

48. Guo C, Ding G, Huang W, Wang Z, Meng Z, Xiao W. Total saponin of Dioscoreae hypoglaucae rhizoma ameliorates streptozotocin-induced diabetic nephropathy. Drug Des Devel Ther 2016; 10: 799–810.

49. Miao Y, Wu X, Xue X, Ma X, Yang L, Zeng X, et al. Morin, the PPARγ agonist, inhibits Th17 differentiation by limiting fatty acid synthesis in collagen-induced arthritis. Cell Biol Toxicol 2023; 39(4): 1433–52.

50. Wen S, He L, Zhong Z, Zhao R, Weng S, Mi H, Liu F. Stigmasterol restores the balance of Treg/Th17 cells by activating the butyrate-PPARγ axis in colitis. Front Immunol 2021; 12: 741934.

